## Supplement Figure S for "Dynamics of Collagen Oxidation and Cross Linking in Regenerating and Irreversibly Infarcted Myocardium"

### Experimental information

#### Synthesis and characterization of compounds

##### Materials and instruments:

NMR spectra were recorded on a JEOL ECZ 500R 11.7 T NMR system equipped with a 5 mm broadband probe (^1^H: 499.81 MHz, ^13^C: 125.68 MHz). ^1^H NMR spectral shifts are reported as singlet (s), doublet (d), triplet (t), quartet (q) or multiplet (m). High resolution mass spectra were acquired on a high-resolution time-of-fight mass spectrometer (AccuTOFDART, JEOL). UV-Vis spectra were recorded on a SpectraMax M2 spectrophotometer using cuvettes with a 1 cm path length. MilliQ purified water was used for all reactions and experiments where water is the solvent. 6-carboxytetramethyl rhodamine was purchased from Fisher and used without purification. TAMRA hydrazide, 6 isomer was purchased from Lumiprobe and used without further purifications. Tert-Butyl 3-aminopropoxycarbamate was purchased from AmBeed and used without further purification. All other chemical supplies and reagents were purchased from common commercial suppliers and used without purifications unless otherwise noted.

**HPLC-MS:** LC-MS analysis was carried out on an Agilent 1260 system (UV detection at 220, 254 and 550 nm) coupled to an Agilent Technologies 6130 Quadrupole MS system. Mobile Phases: A: 0.1% formic acid in H_2_O (v/v) B: 0.1% formic acid in CH_3_CN (v/v) C: 10 mM NH_4_OAc in H_2_O, D: 90% CH_3_CN + 10% solvent C. UV detection at 220, 254 and 280 and 550 nm.

Method 1: Column: Phenomenex LUNA, C18(2), 5 µm, 100 × 2 mm, flow rate: 0.7 mL/min. UV-Vis detection, 220, 254, 280 and 550 nm.

| Time (min) | %A | %B |
| --- | --- | --- |
| 0 | 95 | 5 |
| 3 | 5 | 95 |
| 4.5 | 5 | 95 |
| 5 | 95 | 5 |
| 7 | 95 | 5 |

Method 2: Column: Phenomenex LUNA, C18(2), 5 µm, 100 × 2 mm, flow rate: 0.7 mL/min UV-Vis detection, 220, 254, 280 and 550 nm.

| Time (min) | %C | %D |
| --- | --- | --- |
| 0 | 95 | 5 |
| 0.5 | 95 | 5 |
| 7.5 | 5 | 95 |
| 8.5 | 95 | 5 |
| 10 | 95 | 5 |

**Flash chromatography:** Large-scale reverse-phase purifications were carried out on a Teledyne ISCO CombiFlash system with UV-Vis detection at 220 and 254 nm.

Method 3: Column: 150 g C18, flow rate: 70 mL/min. Mobile Phases: A: 0.1% formic acid in H_2_O (v/v) B: 0.1% formic acid in CH_3_CN (v/v).

| Time (min) | %A | %B |
| --- | --- | --- |
| 0 | 95 | 5 |
| 5 | 95 | 5 |
| 30 | 30 | 70 |
| 35 | 5 | 95 |
| 40 | 5 | 95 |

Method 4: Column: 150 g Silica, flow rate: 100 mL/min. Mobile Phases: A: Hexanes, B: ethyl acetate.

| Time (min) | %A | %B |
| --- | --- | --- |
| 0 | 95 | 5 |
| 10 | 95 | 5 |
| 24 | 40 | 60 |
| 40 | 0 | 100 |

Method 5: Column: 15.5 g C18, flow rate: 40 mL/min. Mobile Phases: A: 0.1% formic acid in H_2_O (v/v) B: 0.1% formic acid in CH_3_CN (v/v).

| Time (min) | %A | %B |
| --- | --- | --- |
| 0 | 95 | 5 |
| 5 | 95 | 5 |
| 15 | 30 | 70 |
| 17 | 5 | 95 |
| 20 | 5 | 95 |

**Preparative HPLC:** Preparative reversed-phase HPLC with UV detection at 220, 254 and 280 nm was performed using Agilent 1260 system. Mobile Phases: A: 0.1% formic acid in H_2_O (v/v) B: 0.1% formic acid in CH_3_CN (v/v) C: 0.1% trifluoroacetic acid in H_2_O (v/v) D: 0.1% trifluoroacetic acid in CH_3_CN (v/v). UV detection at 220, 254 and 280 nm

Method 6: Column: Phenomenex LUNA C18(2) 10 µm, 250 × 21.2 mm, flow rate: 15 mL/min

| Time (min) | %A | %B |
| --- | --- | --- |
| 0 | 95 | 5 |
| 20 | 55 | 45 |
| 40 | 5 | 95 |
| 48 | 5 | 95 |
| 50 | 95 | 5 |

**Analytical HPLC:** HPLC with fluorescence detection was acquired on an Agilent 1260 with binary pump, autosampler, multi-wavelength detector, thermostatted column compartment, and vacuum degasser and a fluorescence detector.

Method 7: Column, Xbridge, 5μm C18 3.5 mm, 150x4.6 mm, flow rate: 1.0 mL/min

Mobile phases: A: 0.1% trifluoroacetic acid in H_2_O (v/v) B: 0.1% trifluoroacetic acid in CH_3_CN (v/v). UV detection at 220, 254, 280 and 550 nm. Fluorescence detection at excitation 545 nm, emission 566 nm.

| Time (min) | %A | %B |
| --- | --- | --- |
| 0 | 95 | 5 |
| 3 | 5 | 95 |
| 4.5 | 5 | 95 |
| 5 | 95 | 5 |
| 7 | 95 | 5 |

Method 8: Column, TSKgel™ ODS-80Tm HPLC Column, 5 μm Particle Size, 150x4.6 mm, flow rate: 1.0 mL/min. Mobile phases: A: 0.15% HFBA, 24% Methanol in H_2_O (v/v) B: 0.1% HFBA, 40% Methanol in H_2_O (v/v) , C: CH_3_CN. UV detection at 220, 254 and 295 nm. Fluorescence detection at excitation 295 nm, emission 400 nm.

| Time (min) | %A | %B | %C |
| --- | --- | --- | --- |
| 0 | 100 | 0 | 0 |
| 17 | 100 | 0 | 0 |
| 17.1 | 0 | 100 | 0 |
| 30 | 0 | 100 | 0 |
| 30.1 | 25 | 0 | 75 |
| 35 | 25 | 0 | 75 |
| 35.1 | 100 | 0 | 0 |
| 40 | 100 | 0 | 0 |

##### *Synthetic schemes and procedure:*

##### Synthesis of TMR-O:

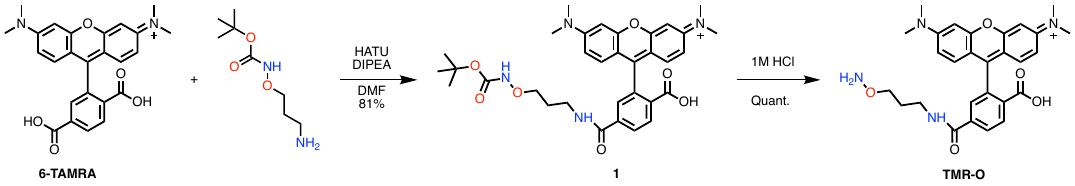

**Compound 1-1**. To a solution of 6-carboxy tetramethylrhodamine (6-TAMRA, 22 mg, 51 μmol) and HATU (19 mg, 51 μmol) in DMF (1 mL) was added  tert-Butyl 3-aminopropoxycarbamate (9.7 mg, 51 μmol) in DMF (0.1 mL). The mixture was stirred for 45 min, then purified by Prep-HPLC, Method 6 using A and B as eluents. . Compound 1-1 was isolated as a red solid after freeze-drying. Mass= 25 mg, 81%. LC-MS (Method 1): t_R_= 3.10 min. m/z =603.2 [M+]; calculated for [C_33_H_39_N_4_O_7_]^+^ 603.28. Purity: >99% (LC-MS, 550 nm). ^1^H NMR (500 MHz, Methanol-*d*_4_) δ 8.39 (d, *J* = 8.2 Hz, 1H), 8.25 (dd, *J* = 8.3, 1.8 Hz, 1H), 7.88 (d, *J* = 1.8 Hz, 1H), 7.16 – 7.13 (m, 2H), 7.03 (dd, *J* = 9.5, 2.4 Hz, 2H), 6.94 (d, *J* = 2.5 Hz, 2H), 3.89 (t, *J* = 5.8 Hz, 2H), 3.55 (t, *J* = 6.6 Hz, 2H), 3.29 (s, 12H), 1.90 (p, *J* = 6.2 Hz, 2H), 1.39 (s, 9H). ^13^C NMR (126 MHz, D_2_O and Acetonitrile-*D*_3_) δ 167.31, 167.30, 159.45, 158.30, 138.91, 138.75, 134.93, 133.94, 132.65, 131.73, 130.18, 129.22, 128.64, 115.28, 114.21, 97.18, 82.49, 75.45, 41.22, 38.35, 28.33, 27.90.

**TMR-O**. Compound 1-1 (25 mg, 41μmol) was suspended in aqueous 1M HCl and stirred overnight protected from light. The solvent was then freeze-dried, resulting in TMR-O as a red solid. Mass= 21 mg, quantitative. LC-MS (Method 1): t_R_= 2.55 min, m/z =503.30 [M+]; calculated for [C_28_H_31_N_4_O_5_]^+^: 503.23. Purity: >99% (LC-MS, 550 nm). ^1^H NMR (500 MHz, METHANOL-*D*_4_) δ 8.40 (d, *J* = 8.2 Hz, 1H), 8.20 (dd, *J* = 8.2, 1.8 Hz, 1H), 7.82 (d, *J* = 1.8 Hz, 1H), 7.12 (d, *J* = 9.5 Hz, 2H), 7.04 (dd, *J* = 9.5, 2.5 Hz, 2H), 6.99 (d, *J* = 2.5 Hz, 2H), 4.10 (t, *J* = 6.1 Hz, 2H), 3.50 (t, *J* = 7.0 Hz, 3H), 3.30 (s, 12H), 1.98 (t, *J* = 7.0 Hz, 2H). ^13^C NMR (126 MHz, Methanol-*D*_4_) δ 167.63, 166.82, 159.21, 157.77, 157.67, 140.40, 138.91, 137.83, 134.30, 133.63, 131.55, 130.69, 129.11, 128.64, 127.38, 127.34, 126.00, 114.24, 113.53, 96.15, 72.70, 39.61, 36.32, 27.68.

##### Synthesis of TMR-HZN:

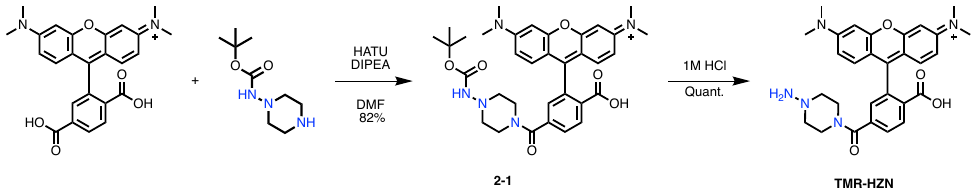

**Compound 2-1**. To a solution of 6-carboxy tetramethylrhodamine (6-TAMRA, 22 mg, 51 μmol) and HATU (19 mg, 51 μmol) in DMF (1 mL) was added Tert-Butyl piperazin-1-ylcarbamate (11.3 mg, 51 μmol) in DMF (0.1 mL). The mixture was stirred for 45 min, then purified by Prep-HPLC, Method 6 using A and B as eluents. . Compound 2-1 was isolated as a red solid after freeze-drying. Mass= 20 mg, 81%. LC-MS (Method 1): t_R_= 2.99 min, m/z =614.30 [M+]; Calculated for [C_34_H_40_N_5_O_6_]+ 614.30. Purity: >99% (LC-MS, 550 nm). ^1^H NMR (500 MHz, D_2_O : CD_3_CN 10:1 ) δ 8.05 (dq, *J* = 4.1, 1.7 Hz, 2H), 7.84 (dd, *J* = 8.5, 1.9 Hz, 1H), 7.09 (dd, *J* = 9.4, 2.1 Hz, 2H), 6.71 (dd, *J* = 9.5, 2.5 Hz, 2H), 6.45 (d, *J* = 2.6 Hz, 2H), 3.00 (d, *J* = 2.2 Hz, 12H), 2.74 (t, *J* = 5.0 Hz, 4H), 2.62 (s, 4H), 1.33 (s, 9H). ^13^C NMR (126 MHz, D_2_O : CD_3_CN 10:1) δ 172.76, 172.38, 164.80, 162.72, 158.38, 155.91, 155.88, 155.56, 141.09, 137.14, 136.75, 129.80, 117.66, 114.52, 112.84, 112.03, 95.30, 43.06, 39.15, 26.83, 26.66.

**TMR-HZN.** Compound 2-1 ( 27 mg, 44 μmol) was suspended in aqueous 1M HCl and stirred overnight protected from light. The solvent was then freeze-dried, resulting in compound 21 as a red solid. Mass= 23 mg, quantitative. LC-MS (Method 1): t_R_= 3.0 min, m/z =514.2 [M+]; Calculated for [C_29_H_32_N_5_O_4_]^+^ 514.24. ^1^H NMR (500 MHz, Acetonitrile-*D*_3_) δ 8.30 (d, *J* = 8.1 Hz, 1H), 7.74 (dd, *J* = 8.1, 1.7 Hz, 1H), 7.35 (d, *J* = 1.7 Hz, 1H), 7.13 (d, *J* = 9.4 Hz, 2H), 6.91 (dd, *J* = 9.4, 2.4 Hz, 2H), 6.81 (d, *J* = 2.4 Hz, 2H), 3.63 (m, 2H), 3.22 (s, 12H), 3.07 (m, 2H). ^13^C NMR (126 MHz, D_2_O : CD_3_CN 10:1) δ 172.86, 172.92, 164.95, 163.27, 155.95, 155.91, 155.62, 141.59, 138.24, 137.15, 130.20, 117.71, 114.30, 111.48, 111.29, 95.20, 62.10, 47.59, 39.52.

##### Synthesis of TMR-Pyr:

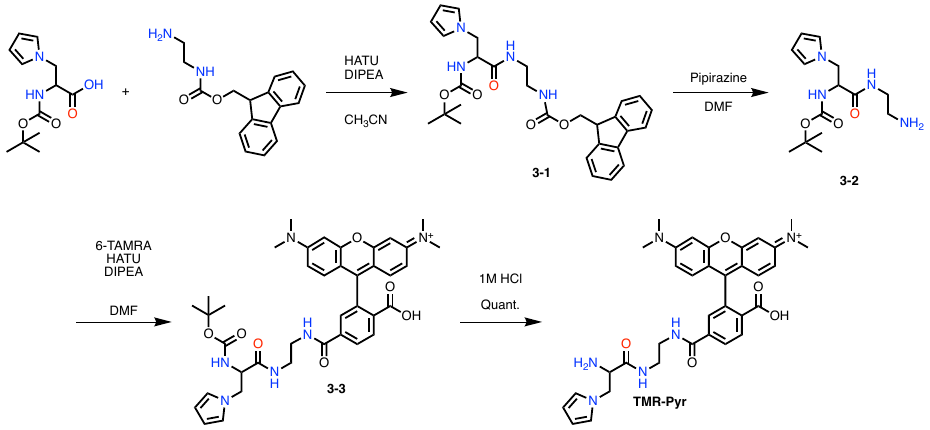

**Compound 3-1**. To a solution of 2-{(tert-butoxy)carbonyl]amino}-3-(1H-pyrrol-1-yl)propanoic acid (100 mg, 0.39 mmol), and N-Fmoc-ethylenediamine hydrochloride (126 mg, 0.39 mmol) in anhydrous DMF (1 mL) was added HATU (151 mg, 0.39 mmol) and diisopropyl ethylamine (DIPEA, 180 μL, 1.0 mmol). The mixture was stirred for one hour, with monitoring by LC-MS. After 3 hours, product was precipitates by addition of water, and then purified by combiflash using Method 5 . Product was isolated as white solid after freeze-drying. Mass = 80 mg, 40% yield. LC-MS (Method 1): t_R_= 4.13 min, m/z = 519.2 [M+H]^+^ ; Calculated for [C_29_H_35_N_4_O_5_]^+^ 519.26. ^1^H NMR (500 MHz, Chloroform-*d*) δ 7.79 (dd, *J* = 7.6, 3.6 Hz, 2H), 7.59 (d, *J* = 7.5 Hz, 2H), 7.41 (t, *J* = 7.5 Hz, 2H), 7.38 – 7.28 (m, 2H), 6.60 (t, *J* = 2.1 Hz, 2H), 6.04 – 5.97 (m, 2H), 5.75 (s, 1H), 5.16 – 5.07 (m, 1H), 4.85 (t, *J* = 6.2 Hz, 1H), 4.46 (ddd, *J* = 48.0, 10.6, 6.3 Hz, 2H), 4.37 – 4.15 (m, 3H), 4.00 – 3.91 (m, 1H), 3.37 – 3.11 (m, 2H), 3.11 – 2.90 (m, 2H), 1.44 (s, 9H). ^13^C NMR (126 MHz, Chloroform-*D*) δ 170.02, 170.02, 156.76, 155.19, 151.73, 143.94, 127.83, 127.20, 125.19, 121.25, 120.07, 109.20, 66.31, 51.76, 51.51, 51.01, 47.50, 44.77, 40.97, 39.99, 28.38.

**Compound 3-2**. To a solution of compound 3-1 ( 80 mg, 0.15 mmoles) in DMF (1 mL) was added piperazine (100 mg, 1.16 mmol) and the reaction was stirred at room temperature for 3 hours. The crude mixture was purified by CombiFlash using Method 5 to result in 3-2 as a white solid, mass =39 mg, 87 % yield. LC-MS (Method 1): t_R_= 2.89 min, m/z = [[M+H]^+^ ; Calculated for [C_14_H_25_N_4_O_3_]^+^ 297.18 no mass visible by LC-MS. ^1^H NMR (500 MHz, D2O, formic acid reference) 6.64 (q, *J* = 2.0 Hz, 2H), 6.05 (q, *J* = 2.0 Hz, 2H), 4.29 (ddd, *J* = 7.8, 5.8, 1.7 Hz, 1H), 4.23 – 4.03 (m, 2H), 3.45 – 3.23 (m, 2H), 2.93 (q, *J* = 9.4, 7.9 Hz, 2H), 1.26 (s, 9H). ^13^C NMR (126 MHz, D_2_O) δ 168.15, 166.00, 152.34, 145.73, 116.96, 103.56, 103.56, 77.03, 51.11, 44.58, 33.98, 31.87, 22.60.

**Compound 3-3**.To a solution of TAMRA (17 mg, 39 μmol) and HATU (16 mg, 43 μmol) in DMF (0.5 mL) was added dropwise a solution of **3-2** (17 mg, 58 μmol) in DMF (0.1 mL). The mixture was stirred for 2 hrs, then purified by Prep-HPLC, Method 6 using A and B as eluents. **Compound 3-3** was isolated as a red solid after freeze-drying. Mass= 23 mg, 83%. LC-MS (Method 1): t_R_= 4.3 min, m/z =709.3 [M+]; Calculated for [C_39_H_45_N_6_O_7_]^+^ 709.33. >95.5% purity (LC-MS, 550 nm). ^1^H NMR (500 MHz, Acetonitrile-*D*_3_) δ 8.28 (d, *J* = 8.2 Hz, 1H), 8.15 (d, *J* = 8.2 Hz, 1H), 7.86 (s, 2H), 7.29 (s, 1H), 7.04 (dd, *J* = 9.5, 3.5 Hz, 2H), 6.81 (ddd, *J* = 9.5, 3.5, 2.1 Hz, 2H), 6.78 – 6.68 (m, 2H), 6.40 (s, 2H), 5.92 (t, *J* = 2.1 Hz, 2H), 5.82 (d, *J* = 8.0 Hz, 1H), 4.11 – 4.00 (m, 2H), 3.78 (dd, *J* = 15.0, 9.7 Hz, 1H), 3.48 (s, 1H), 3.18 (s, 14H), 2.97 (td, *J* = 6.6, 3.8 Hz, 2H), 1.72 – 1.67 (m, 2H), 1.22 (s, 9H).^13^C NMR (126 MHz, Methanol-*D*_4_) δ 171.61, 168.65, 166.51, 166.01, 159.30, 157.65, 140.31, 137.89, 134.25, 133.39, 131.53, 130.66, 128.80, 128.13, 120.72, 114.24, 113.50, 107.89, 96.12, 79.68, 56.22, 48.19, 48.02, 47.85, 47.68, 47.51, 47.34, 47.16, 39.61, 38.48, 27.20, 26.13, 24.01.

**TMR-Pyr.** Compound **3-3** was suspended in aqueous 1M HCl and stirred overnight protected from light. The solvent was then freeze-dried, resulting in compound 21 as a red solid. Mass= 20 mg, quantitative. LC-MS (Method 1): t_R_= 2.58 min, m/z =609.2 [M+]; Calculated for [C_34_H_37_N_6_O_5_]^+^ 709.33609.28. >99.5% purity (LC-MS, 550 nm). ^1^H NMR (500 MHz, Acetonitrile-*D*_3_) δ 8.28 (d, *J* = 8.2 Hz, 1H), 8.15 (d, *J* = 8.2 Hz, 1H), 7.86 (s, 2H), 7.29 (s, 1H), 7.04 (dd, *J* = 9.5, 3.5 Hz, 2H), 6.81 (ddd, *J* = 9.5, 3.5, 2.1 Hz, 2H), 6.78 – 6.68 (m, 2H), 6.40 (s, 2H), 5.92 (t, *J* = 2.1 Hz, 2H), 5.82 (d, *J* = 8.0 Hz, 1H), 4.09 – 4.00 (m, 2H), 3.78 (dd, *J* = 15.0, 9.7 Hz, 1H), 3.46 (d, *J* = 22.2 Hz, 2H), 3.21 (s, 2H), 3.16 (d, *J* = 18.9 Hz, 13H), 2.97 (td, *J* = 6.6, 3.8 Hz, 2H), 2.06 – 2.03 (m, 1H), 1.72 – 1.67 (m, 2H).^13^C NMR (126 MHz, Acetonitrile-*D*_3_) δ 186.88, 171.37, 167.19, 166.68, 166.20, 160.54, 158.15, 138.99, 134.10, 132.28, 131.83, 131.76, 129.83, 121.90, 118.26, 115.00, 114.16, 108.77, 97.13, 50.21, 46.89, 41.20, 28.32, 26.95, 1.74, 1.58, 1.41, 1.25, 1.08, 0.92, 0.75.

##### Synthesis of TMR-Rho

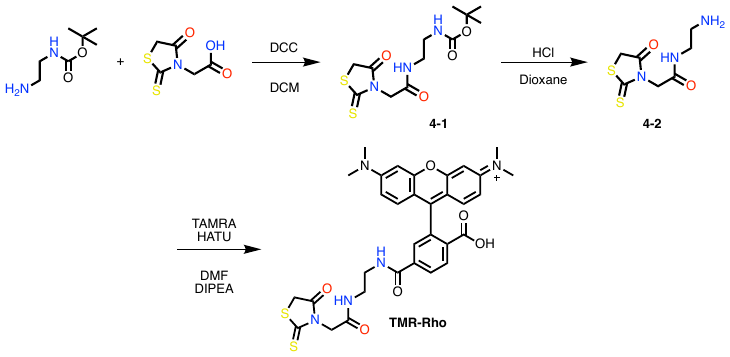

**Compound 4-1**. To a solution of rhodanine-3-carboxylic acid (1g, 5.2 mmol) in dichloromethane (20 mL) was added DCC (1.6 g, 7.8 mmol), and N-hydroxy succinimide (800 mg, 7 mmol) and stirred for 5 minutes, resulting in an organge solution. To this orange solution was added mono-Fmoc ethylene diamine hydrochloride (1.9 g, 6.2 mmole) in DMF (10 mL), and the combined solution was stirred for an additional 10 hours. The resulting dark orange solution was purified by Combiflash using Method 5. The product was isolated as an orange oil, mass= 0.8 g, 47% yield. ^1^H NMR (500 MHz, Acetonitrile-*d*_3_) δ 6.94 (s, 1H), 5.45 (s, 1H), 4.51 (s, 2H), 4.13 (s, 2H), 3.21 (q, *J* = 6.0 Hz, 2H), 3.09 (t, *J* = 6.1 Hz, 2H), 1.40 (s, 9H). ^13^C NMR (126 MHz, Acetonitrile-*d*_3_) δ 203.39, 174.11, 165.56, 156.43, 117.47, 78.65, 46.33, 39.83, 39.57, 36.04, 27.71

**Compound 4-2**. Compound 4-1 (0.8 g, 2.4 mmol) was placed in 1M HCl in dioxane and stirred overnight. Solvent was removed by freeze-drying, yielding compound 4-2 as an orange powder which was used without further purification. Mass product 0.56 g, quantitative yield. ^1^H NMR (500 MHz, DMSO-*d*_6_) δ 8.57 (t, *J* = 5.7 Hz, 1H), 8.05 (s, 2H), 4.51 (s, 2H), 4.34 (s, 2H), 3.31 (q, *J* = 6.3 Hz, 2H), 2.83 (h, *J* = 6.1 Hz, 2H). ^13^C NMR (126 MHz, DMSO-*D*_6_) δ 203.68, 174.52, 165.87, 46.55, 38.74, 37.04, 36.71.

**TMR-Rho**. To a solution of 6-TAMRA (50 mg, 116 μmol) and HATU (46 mg, 122 μmol) in DMF (0.5 mL) was added **compound 4-2** (30 mg, 0.13 μmol) in DMF (0.1 mL). The mixture was stirred for 2 hrs, then purified by Combiflash, Method 5 using A and B as eluents. TMR-Rho was isolated as a red solid after freeze-drying. Mass= 45 mg, 62%. LC-MS (Method 1): t_R_= 2.89 min, m/z =646.0 [M+]; Calculated for [C_32_H_32_N_5_O_6_S_2_]^+^ 646.18. Purity = >99.0% (LCMS, 550nm). ^1^H NMR (500 MHz, Methanol-*D*_4_) δ 8.30 (d, *J* = 8.2 Hz, 1H), 8.11 (dd, *J* = 8.2, 1.8 Hz, 1H), 7.73 (d, *J* = 1.8 Hz, 1H), 7.18 (d, *J* = 9.5 Hz, 2H), 7.03 (dd, *J* = 9.5, 2.5 Hz, 2H), 6.96 (d, *J* = 2.5 Hz, 2H), 4.54 (s, 2H), 4.08 (s, 2H), 3.53 – 3.46 (dd, , *J* = 4.6, 2.7 Hz, 2H), 3.43 (dd, *J* = 4.6, 2.7 Hz, 2H), 2.63 (s, 12H). ^13^C NMR (126 MHz, Methanol-*D*_4_) δ 194.87, 185.39, 179.93, 168.63, 167.26, 167.19, 159.15, 158.94, 155.91, 152.76, 151.72, 135.12, 132.38, 130.07, 130.07, 115.46, 115.02, 97.41, 78.00, 49.51, 49.34, 49.17, 49.00, 48.83, 48.66, 48.49, 40.92, 40.39, 40.39.

Synthesis of TMR-LHZ
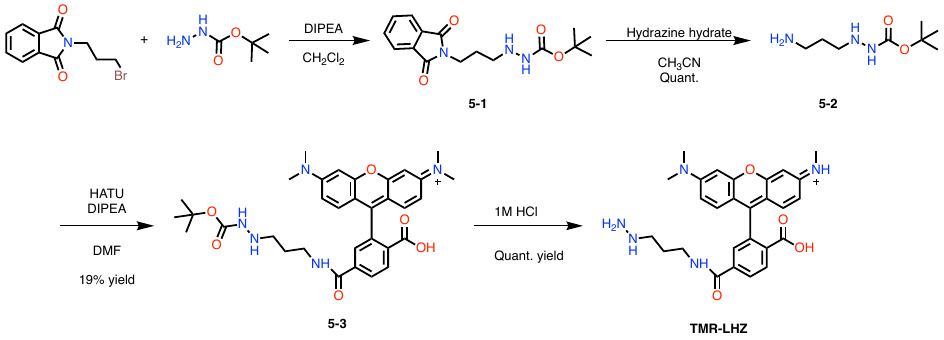

**Compound 5-1**. To a solution of N-(4-Bromobutoxy)phthalimide (5 g, 18.7 mmol) in dichloromethane (50 mL) was added a solution of t-butylcarbazate (4.9 g, 37.3 mmol) in 10 mL dichloromethane containing DIPEA (3.9 mL, 22.4 mmol). The resulting clear solution was stirred overnight then purified by Combiflash using silica column and Method 4. The resulting product was isolated as a clear oil that solidified on standing. Mass product = 4 g, 67% yield. ^1^H NMR (500 MHz, Chloroform-*d*) δ 7.80 (dtd, *J* = 7.9, 4.8, 2.3 Hz, 2H), 7.68 (pd, *J* = 6.6, 5.7, 3.3 Hz, 2H), 6.24 (s, 1H), 3.75 (q, *J* = 6.7 Hz, 2H), 2.87 (q, *J* = 6.7 Hz, 2H), 1.81 (dp, *J* = 6.5 Hz, 2H), 1.41 (s, Hz, 9H). ^13^C NMR (126 MHz, Chloroform-*d*) δ 168.53, 156.92, 133.99, 132.22, 123.29, 80.55, 49.20, 35.87, 28.41, 26.82.

**Compound 5-2**. To a solution of compound 5-1 (1.9 g, 0.62 mmol) in acetonitrile (50 mL) was added hydrazine hydrate (1 mL, 2 mmol) and the solution was brought to reflux and heated overnight. The resulting mixture was filtered through celite plug, and the clear solution was evaporated to dryness via rotary evaporation. The resulting clear oil was dried under high vacuum overnight and used without further purification. Mass, 1.1 g, quantitative yield. ^1^H NMR (500 MHz, DMSO-*d*_6_) δ 9.36 (s, 1H), 8.28 (s, 1H), 7.68 (s, 2H), 2.83 (s, 2H), 2.71 (t, *J* = 7.0 Hz, 2H), 1.57 (p, *J* = 9.2, 8.2 Hz, 2H), 1.36 (s, 9H). ^13^C NMR (126 MHz, DMSO-*d*_6_) δ 156.91, 80.47, 56.21, 50.06, 39.96, 31.30, 28.41.

**Compound 5-3**. To a solution of 6-carboxy tetramethylrhodamine (6-TAMRA, 50 mg, 116 μmol) and HATU (46 mg, 121 μmol) in DMF (1 mL) was added  Compound 5-2 (25 mg, 128 μmol) in DMF (0.1 mL). The mixture was stirred for 1 hr min, then purified by Prep-HPLC, Method 14 using A and B as eluents. Compound 5-3 was isolated as a red solid after freeze-drying. Mass= 13 mg, 19%. LC-MS (Method 1): t_R_= 4.27min, m/z =602.30 [M+]; Calculated for [C_39_H_45_N_6_O_7_]^+^ 709.33602.30. Purity = >99.9% (LCMS, 550nm).

^1^H NMR (500 MHz, METHANOL-*D*_4_) δ 8.38 (d, *J* = 8.2 Hz, 1H), 8.20 (dd, *J* = 8.2, 1.8 Hz, 1H), 7.85 (d, *J* = 1.8 Hz, 1H), 7.12 (d, *J* = 9.5 Hz, 2H), 7.03 (dd, *J* = 9.5, 2.5 Hz, 2H), 6.95 (d, *J* = 2.5 Hz, 2H), 4.89 (s, 12H), 3.47 (t, *J* = 6.6 Hz, 2H), 3.05 (t, *J* = 7.2 Hz, 2H), 1.86 (t, *J* = 6.9 Hz, 2H), 1.41 (s, 10H). ^13^C NMR (126 MHz, Methanol-*D*_4_) δ 166.88, 166.04, 159.28, 157.75, 157.63, 137.80, 134.22, 133.64, 131.50, 130.73, 128.99, 128.78, 114.23, 113.54, 96.14, 81.39, 39.61, 37.50, 27.14, 25.28.

**TMR-LHZ**. Compound 5-3 (13 mg, 22 μmol) was suspended in aqueous 1M HCl and stirred overnight protected from light. The solvent was then freeze-dried, resulting in TMR-LHZ as a red solid. Mass= 21 mg, quantitative yield. LC-MS (Method 1): t_R_= 3.3 min, m/z =502.2 [M+]; calcd: 502.24. Purity = >99.9% (LCMS, 550 nm). ^1^H NMR (500 MHz, Methanol-*D*_4_) δ 8.48 – 8.32 (m, 1H), 8.29 – 8.14 (m, 1H), 7.84 (dd, *J* = 5.2, 2.7 Hz, 1H), 7.19 – 7.09 (m, 2H), 7.05 (tt, *J* = 7.0, 2.9 Hz, 2H), 6.97 (dt, *J* = 5.1, 2.6 Hz, 2H), 3.49 (t, *J* = 7.3 Hz, 2H), 3.29 (d, *J* = 3.2 Hz, 12H), 3.08 (q, *J* = 6.4 Hz, 2H), 1.94 (q, *J* = 7.1 Hz, 2H). ^13^C NMR (126 MHz, Methanol-*D*_4_) δ 167.06, 165.94, 159.20, 157.77, 157.66, 137.68, 134.29, 133.65, 131.56, 130.72, 129.02, 128.76, 114.27, 113.55, 96.15, 39.63, 36.78, 25.38.

##### Synthesis of TMR-NB

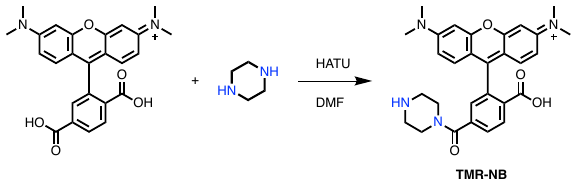

**TMR-NB**. To a solution of TAMRA (10 mg, 23 μmol) and HATU (9.3 mg, 23 μmol) in DMF (0.5 mL) was added piperazine (2.2 mg, 25 μmol) in DMF (0.1 mL). The mixture was stirred for 2 hrs, then purified by Prep-HPLC, Method 14 using A and B as eluents. . Compound 23 was isolated as a red solid after freeze-drying. Mass= 11 mg, quantitative yield. LC-MS (Method 1): t_R_= 2.48 min, m/z =499.2 [M+]; calcd: 499.23. > 99% Purity (LC-MS. 550 nm). ^1^H NMR (500 MHz, Methanol-*D*_4_) δ 8.45 (dd, J= 8.2, 0.7 Hz, 1H), 8.18 (dd, J= 8.2, 1H) 0.7 Hz (m, 1H), 7.94 (dd, *J* = 2.0, 0.7 Hz, 1H), 7.16 – 7.1 (m, 2H), 7.03 (dd, *J* = 9.5, 2.9 Hz, 2H), 6.94 (d, *J* = 2.6 Hz, 2H), 3.32 (t, *J* = 7.2 Hz, 4H), 3.28 (s, 12H), 2.70 (t, *J* = 7.1 Hz, 4H). ^13^C NMR (126 MHz, Methanol-*D*_4_) δ 166.1, 165.9, 159.5, 157.8, 157.7, 134.68, 134.39, 134.05, 131.25, 130.83, 130.0, 129.76, 114.32, 113.65, 96.15, 39.53, 52.82, 47.42.

### Supplementary Figures

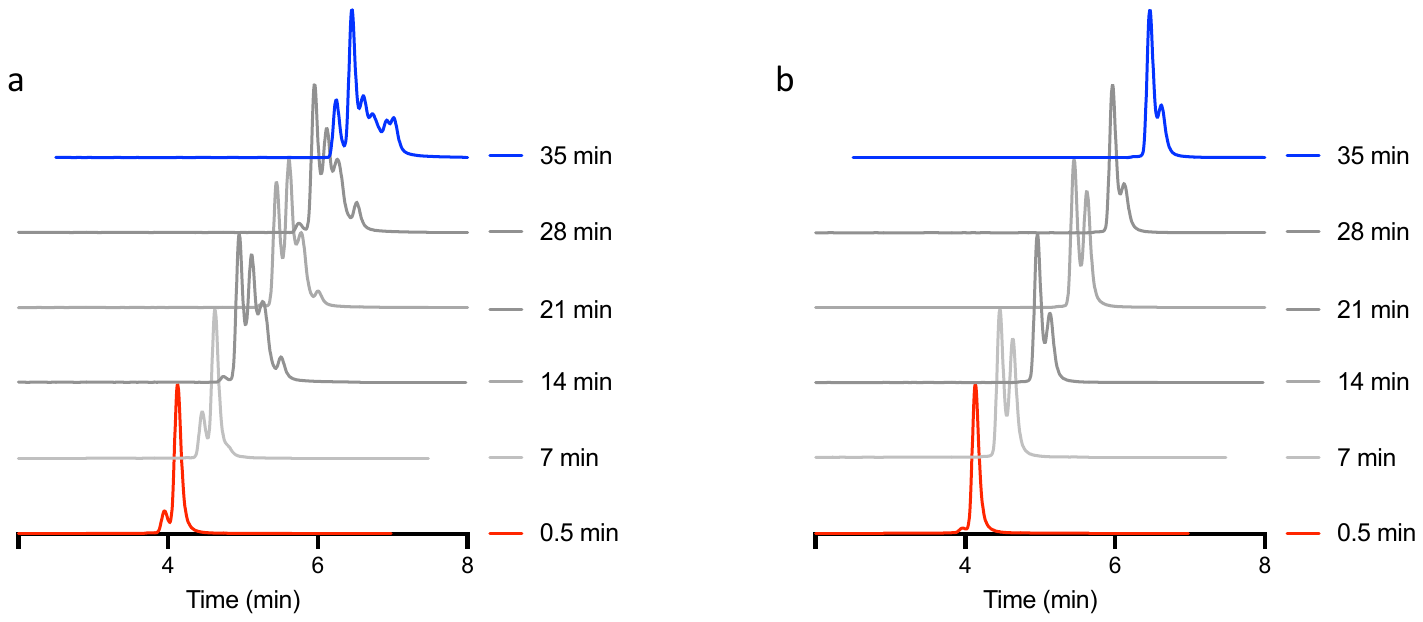

Figure S 1: HPLC traces of TMR-Rho reactions. a) HPLC traces of the reaction of TMR-Rho with butyraldehyde detected by fluorescence (excitation 545 nm, emission 566 nm) showing multiple reaction products. b) HPLC traces of the reaction of TMR-Rho with ethanolamine detected by fluorescence (excitation 545 nm, emission 566 nm) showing a single reaction product. Reaction conditions(0.4 mL pH 7.40 PBS, 10 µM and 110 µM butyraldehyde or ethanolamine.

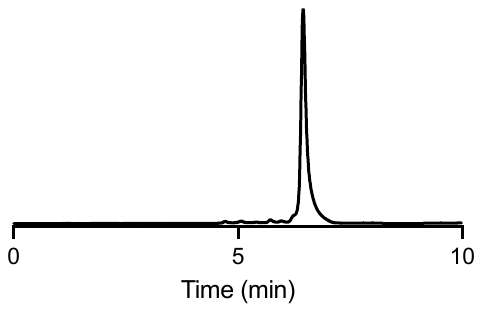

Figure S 2: HPLC traces of TMR-O-butyraldehyde adduct after 24 hours. HPLC trace of TMR-O-butyraldehyde oxime after 24 hours showing little to no hydrolysis observed. Fluorescence detection with excitation at 545 nm, and emission 566 nm.
